# Machine learning-based framework for predicting human infection potential of coronavirus associated with tri-Amino acid motifs, KIQ and LEP in spike protein

**DOI:** 10.64898/2026.02.02.703238

**Authors:** Narathit Chanraeng, Junwen Guo, Tarapong Srisongkram, Yothin Hinwan, Peter Fransson, Henrik Sjödin, Yoshiharu Matsuura, Hans Jorgen Overgaard, Watcharapong Panthong, Tipaya Ekalaksananan, Chamsai Pientong, Supranee Phanthanawiboon

## Abstract

Assessing the human infection potential of emerging coronaviruses remains a critical challenge for global health preparedness. In this study, we developed a machine learning-based framework to predict the human infection potential of coronaviruses and to identify associated sequence motifs using spike (S) protein sequences. A total of 3,904 complete S protein sequences were collected, annotated as human or non-human infection and encoded using trimer-based k-mer features. Model benchmarking was conducted across 27 machine learning algorithms, followed by hyperparameter optimization of the selected model. Robustness and generalizability were evaluated using k-fold cross-validation and independent external validation. Feature interpretability was further assessed using SHAP analysis to identify sequence determinants associated with infection potential. The Random Forest classifier achieved the best performance, with accuracy, sensitivity, and specificity of 97.8%, 99%, and 97.4%, respectively, and demonstrated stable predictive performance across validation datasets. Notably, the KIQ and LEP motifs were strongly associated with human infection coronaviruses and mapped to the HR1 and N-terminal domain regions of the S protein. Overall, this framework provides a practical approach for risk assessment and surveillance of emerging coronaviruses.

**Author summary:** Emerging coronaviruses continue to threaten global public health, but rapidly identifying viruses with the potential to infect humans remains challenging. Traditional experimental approaches are time-consuming and resource-intensive, limiting their use for large-scale surveillance. In this study, we developed a machine learning based workflow to assess the human infection potential of coronaviruses using spike protein sequences. By analyzing sequence patterns across a diverse set of coronaviruses, our framework enables rapid screening of coronaviruses from multiple host species. Unlike previous studies focused on limited coronavirus genera, our approach integrates all four genera and systematically evaluates multiple learning strategies. Importantly, our analysis identifies conserved sequence motifs linked to human infection potential, bridging predictive performance with biological interpretability. Our findings demonstrate computational approaches support early warning systems for identifying high risk coronaviruses, contributing to prioritize viruses for experimental validation, guide surveillance efforts, and strengthen global pandemic preparedness under a One Health perspective.

## Introduction

Coronaviruses have been associated with human emergence. They are subdivided into four genera including genus *Alphacoronavirus* and *Betacoronavirus* exclusively infect mammals, while *Gammacoronavirus* and *Deltacoronavirus* have a wider range of hosts (1). Three major outbreaks of coronaviruses: Severe Acute Respiratory Syndrome Coronavirus (SARS-CoV) in 2002, Middle East Respiratory Syndrome Coronavirus (MERS-CoV) in 2012, and most recently Severe Acute Respiratory Syndrome Coronavirus 2 (SARS-CoV-2), which caused the Coronavirus Disease 2019 (COVID-19) pandemic beginning in 2019 (2–4) have resulted in significant morbidity, mortality, and global economic impact. Furthermore, there are evidence of novel coronavirus can cause emerging event through zoonotic transmission (5, 6). Such emergence events are influenced by factors including viral genetic mutation, human behavior, ecological, climatic change, and host receptor binding specificity (7). Given the severity of coronavirus-associated diseases, predicting human spillover potential of the virus in advance may prevent the experienced effect of the outbreak. Assessing the human infection potential of the existing and newly found coronavirus remains a major challenge.

Coronaviruses are enveloped, single-strand RNA viruses, which cause infection in the respiratory and gastrointestinal tract. To infect host, the virus utilizes the spike (S) glycoprotein attached to the host cells via specific receptors and internalized into the host cell through endocytosis or membrane fusion (8, 9). This protein plays a crucial role in the initial steps of coronavirus infection, making it a crucial target for assessing and predicting infection potential. Receptor binding and entry assessments serve as experimental gold standard approaches for assessing coronavirus infection potential. For example, receptor-binding assays, pseudovirus entry assays, and live virus infection experiments are widely used to evaluate and confirm the ability of viruses to infect human, particularly focusing on the S protein of coronavirus (9–12). However, it is limited by biosafety constraints, scalability, model relevance, and resource demands. To overcome these limitations, computational prediction approaches have been increasingly applied including molecular docking, dynamics simulations, and machine learning (ML) approaches (13–20). Machine learning is a popular and valuable approach for a large-scale screening across multiple virus families. Several studies have developed predictive models by using different techniques to estimate the human infection potential of zoonotic viruses, but technique to predict the performance of coronaviruses remains low accuracy (15, 21, 22). Recent research has attempted to develop a model for predicting human infection and cross species infection potential of coronavirus. However, these studies have focused on alpha-cov and beta-cov but there is limited work on gamma-cov and delta-cov (23, 24).

In this study, we focused on all four genera of coronaviruses, which are known or suspected spillover potential to humans and a wide range of animal hosts. Recognizing this potential, we trained multiple machine learning models using spike protein sequences and host information to evaluate their likelihood of infecting humans. The Random Forest Classifier showed overall strong and consistent performance through cross-validation and independent external testing, indicating that the model can generalize well beyond the training data. In addition, SHAP-based interpretability revealed biologically meaningful tri-amino-acid motifs associated with either human infection or non-human infection viruses. These combination of accuracy and interpretability model can be applied to newly detected or uncharacterized coronaviruses to rapidly estimate their human infection potential, guide laboratory prioritization in in vitro experiments, and support early risk assessment for human infection potential of coronavirus.

## Materials and Methods

### Dataset retrieval and preparing

The amino acid sequences of coronavirus spike proteins were retrieved in full length from the NCBI Virus database on January 21, 2025. To prevent data leakage, duplicate sequences were removed based on sequence identity, resulting in a final dataset of 3,904 unique sequences for model training. Each sequence was annotated as either human infection or non-human infection. Virus sequences isolated from human hosts were labeled as human infection (n = 1,001), while virus sequences isolated from non-human animal hosts were labeled as non-human infection (n = 2,903) (S1 Table). To prepare the dataset, each amino acid sequence was segmented into overlapping trimers (3-mers) by applying a sliding window of size 3 with a step size of 1 amino acid. The resulting trimers were then converted into k-mer features by counting the frequency of each unique trimer within the sequence. The dataset was cleaned by removing any sequences with missing data then randomly split into a training set and a test set with an 80:20 ratio. To address class imbalance in the training set, the Synthetic Minority Over-sampling Technique (SMOTE) was applied to generate synthetic samples for the smaller class (Fig 1A).

**Fig 1.**
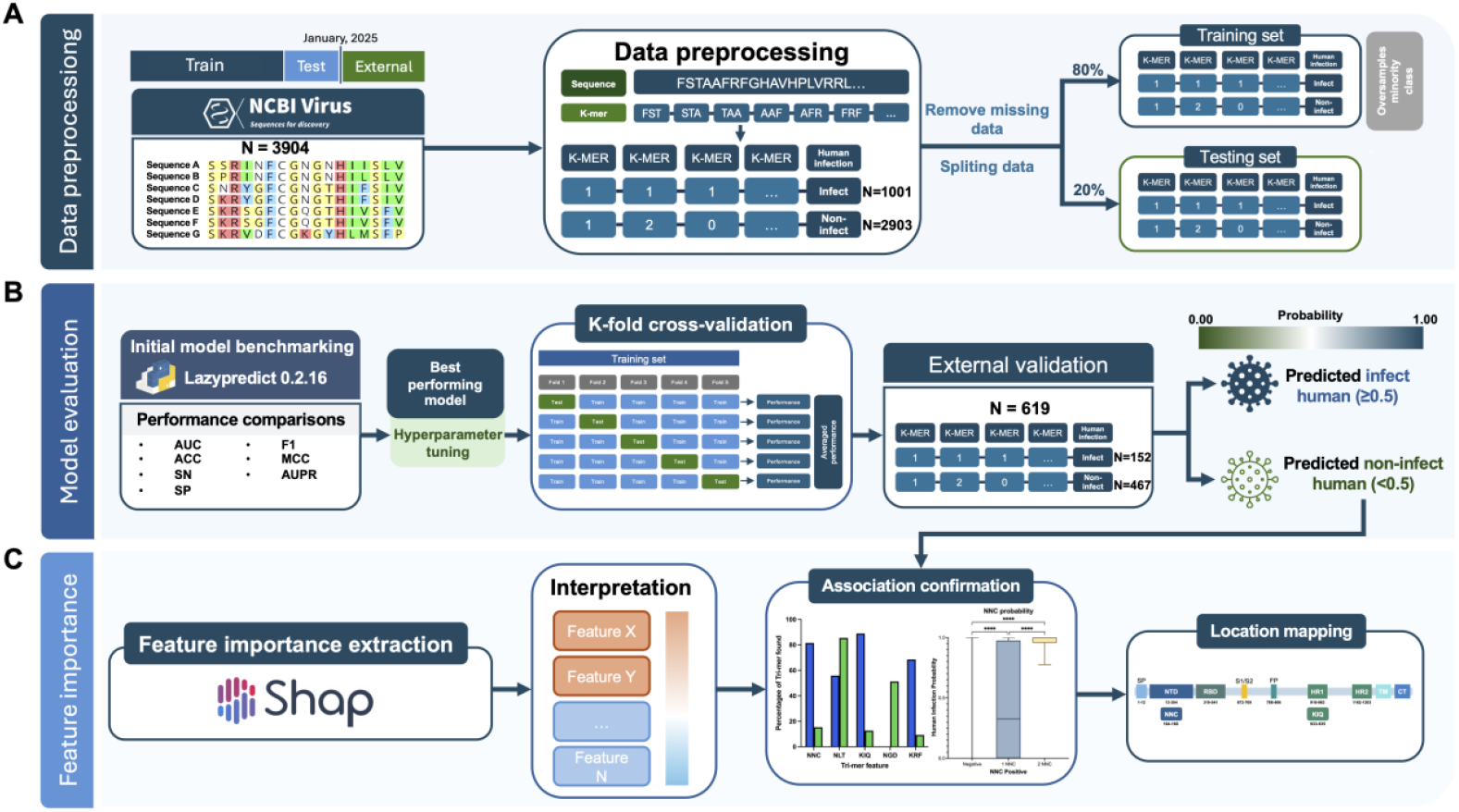
Methodology workflow of machine learning approaches for predicting human infection potential of coronavirus. (A) Data preprocessing: The original and external dataset were retrieved from NCBI virus database in different periods. Then, each sequence was annotated as human infection and non-human infection before transformed into trimer and processing for data training. The original dataset was split into 80:20 for training:testing sets. SMOTE was applied to address class imbalance on training set. (B) Model evaluation: The *Lazypredict* Python package was applied for initial benchmarking to select the best-performing model. Selected model was tuned hyperparameter, then validated by using k-fold cross-validation and external validation. (C) Feature importance: Importance trimers were identified by using SHAP analysis, then confirm association by track back to dataset for biological interpretation and mapped onto each sequence to determine position found.

### Initial model benchmarking

To identify the best-performing model for the dataset, initial benchmarking was conducted using the *Lazypredict* package across 14 repeated runs with different random seeds. *Lazypredict* is a Python package that automates the process of training and evaluating multiple machine learning models with minimal code. It provides quick benchmarking by running several algorithms and reporting their performance metrics, helping to identify the most suitable model for a given dataset. A total of 27 machine learning models spanning diverse algorithmic families were evaluated, including ensemble-based methods, tree-based models, linear models, kernel-based methods, probabilistic classifiers, instance-based methods, discriminant analysis approaches, semi-supervised learning models, as well as baseline and calibration models. Model performance was assessed using key metrics obtained during benchmarking, including accuracy, balanced accuracy, area under the receiver operating characteristic curve (AUC), and F1 score. The average and standard deviation of these metrics across the 14 runs were then computed. Based on these results, the model with the highest balanced performance was chosen for further training and validation (Fig 1B).

### Model hyperparameter tuning and validation

The Random Forest Classifier was selected as the best-performing model for training. Its key hyperparameters, including splitting criterion, the number of trees (n_estimators) and the maximum tree depth (max_depth), were optimized using a grid search with 5-fold cross-validation on the training dataset. The optimal hyperparameter combination was ranked and selected based on the mean AUC across the cross-validation folds. Following hyperparameter tuning, the model was validated using 10-fold cross-validation to assess its predictive performance and generalizability. In this procedure, the dataset was partitioned into 10 subsets (folds), and the process was repeated 10 times so that each fold was used once as the validation set while the remaining folds served as the training set. Model performance was evaluated using accuracy, F1 score, Matthew’s correlation coefficient (MCC), AUC, and area under the precision-recall curve (AUPR), with results reported as mean values ± standard deviation.

### External validation

The external dataset of amino acid sequences of coronavirus spike proteins was retrieved in full length from the NCBI Virus database on June 29, 2025. Sequences were selected based on the criterion that their release date was after January 21, 2025, to ensure they were independent from the training data. Duplicate sequences overlapping with the original training dataset were removed based on sequence identity to prevent data leakage. Each sequence was annotated as either human infection or non-human infection using the same criteria as for the original training dataset. The final external dataset consisted of 619 sequences, including 152 human infection and 467 non-human infection sequences (S2 Table). The final external dataset was used to predict the human infection potential using the developed model, which generated a probability score ranging from 0 to 1.

### Feature importance extraction and confirmation

To identify tri-amino acid motifs potentially associated with human infection, feature importance scores were obtained from the training, test, and external dataset using SHapley Additive exPlanations (SHAP) analysis. SHAP values were calculated to quantify the contribution of each k-mer feature to the predicted probability of human infection. The most important k-mer features, consistently identified across all datasets, were selected as motif and confirmed by mapped back to the complete dataset (including both the original and external datasets), and the percentage occurrence of each motif was compared between the human infection and non-human infection groups. Then, the predicted human infection probability of external dataset predicted by developed model (S2 Table) was plotted against the number of motif occurrences in the dataset to evaluate the association of motif abundance and human infection probability. Finally, the confirmed motifs associated with human and non-human infection were mapped back onto all sequences in the datasets to determine their positional occurrences within each protein sequence (Fig 1C).

### Statistical Analysis

To evaluate the relationship between motif abundance and the Random Forest model-predicted probability of human infection, sequences were grouped by the number of motif occurrences. Differences in predicted probabilities across motif-count groups were assessed using the Kruskal-Wallis test, a non-parametric method that does not assume normality of the data. For motifs showing significant differences, post-hoc pairwise comparisons were performed using Dunn’s test with Bonferroni correction to identify which groups differed significantly. All statistical analyses and visualizations were performed using GraphPad Prism version 9.5.0 (GraphPad Software, Inc., San Diego, CA, USA).

### Declaration of AI-assisted Technology

During preparation of this work ChatGPT-5.2 was used to improve language. After using this tool, the author reviewed and edited the content as needed and take full responsibility for the content of the publication.

## Results

### Comprehensive Benchmarking of ML-Algorithms to predict Human Infection Potential of Coronavirus

The initial benchmarking of candidate classification models was performed to identify the best-performing model for predicting the human infection potential of coronaviruses based on spike protein sequences. Among 27 models tested, the Random Forest Classifier achieved the highest performance, with an accuracy of 0.977, balanced accuracy of 0.981, AUC of 0.981, and an F1 score of 0.977. The SGD Classifier and Logistic Regression models produced identical scores to the Random Forest, indicating similarly strong performance (Fig 2A and S3 Table). To ensure robust results, the initial benchmarking was repeated 14 times using different random seeds. Across these repeated runs, the Random Forest Classifier consistently achieved the highest performance (Fig 2B and S4 Table). Based on its top-ranked performance across all metrics, the Random Forest Classifier was selected for hyperparameter tuning and final model development.

**Fig 2.**
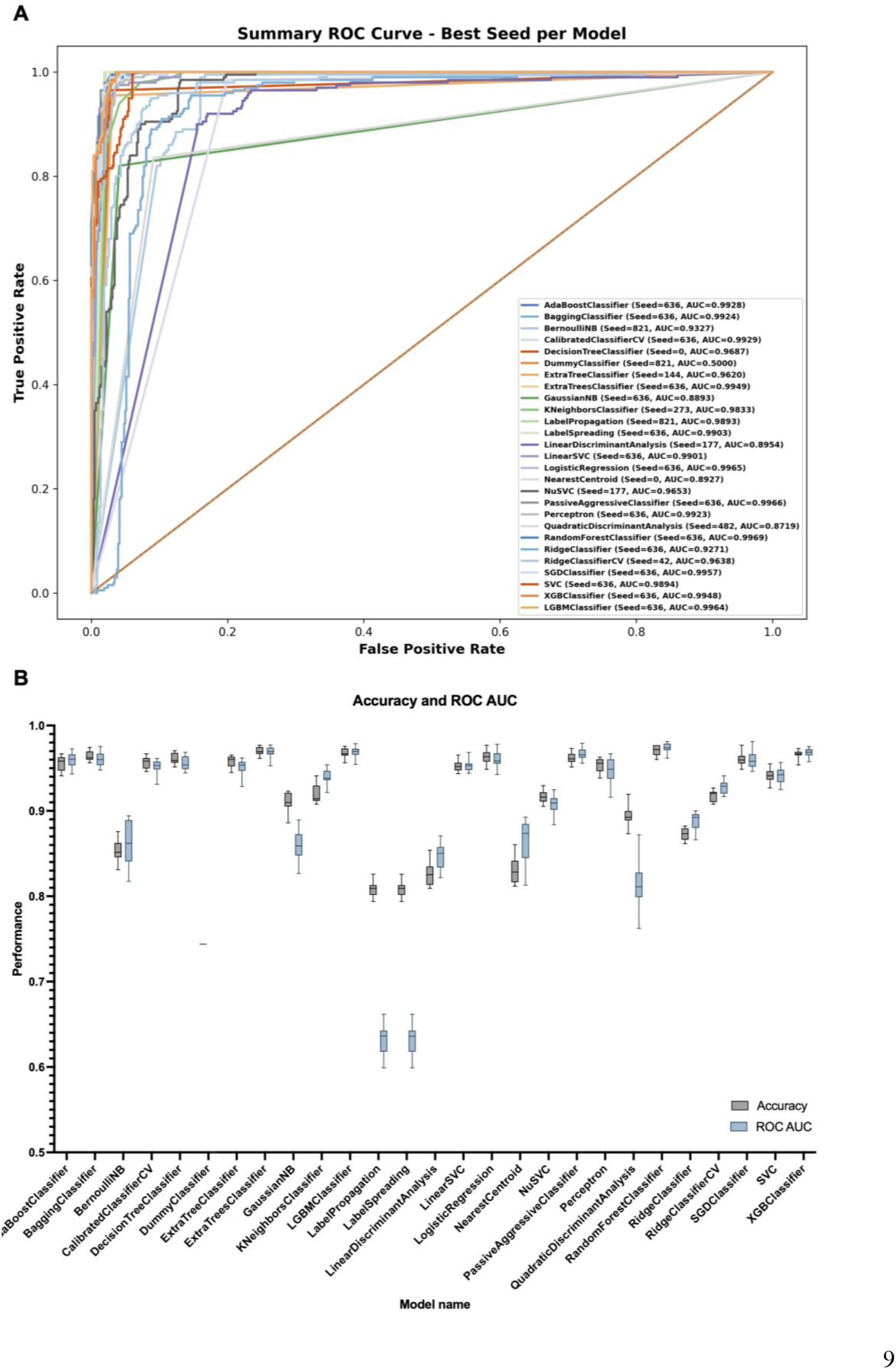
Performance comparison of machine learning models for predicting the human infection potential of coronaviruses. (A) ROC curves of multiple machine learning classifiers evaluated on the test dataset, showing the best-performing random seed for each model. The AUC is indicated in the legend for each classifier. (B) Distribution of model performance metrics across 14 runs, shown as boxplots. Boxes indicate the interquartile range, center lines represent mean values, and whiskers denote the full range of observed values.

### Optimized Random Forest Classifier Demonstrates Excellent Predictive Performance

The Random Forest Classifier was optimized hyperparameters on splitting criterion, maximum tree depth, and number of trees (n_estimators). The best performance has been shown with ‘entropy’, 57, and 210 for those three parameters respectively (S5 Table). The model achieved a sensitivity of 0.9900, indicating a high ability to correctly identify sequences potentially associated with human infection. The specificity was 0.9742, reflecting its effectiveness in correctly classifying non-human infection sequences. The overall accuracy was 0.9782, and the AUC was 0.9969, confirming excellent overall classification performance. The F1 score was 0.9588, indicating a good balance between precision and recall. The AUPR was 0.9905, highlighting the model’s robustness, particularly in handling data imbalance. MCC was 0.9449, further demonstrating the model’s strong predictive power and balanced performance across infection and non-infection group. (Fig 3) These results confirmed that the Random Forest Classifier provides reliable and accurate predictions for predicting human infection potential based on coronavirus spike protein sequences.

**Fig 3.**
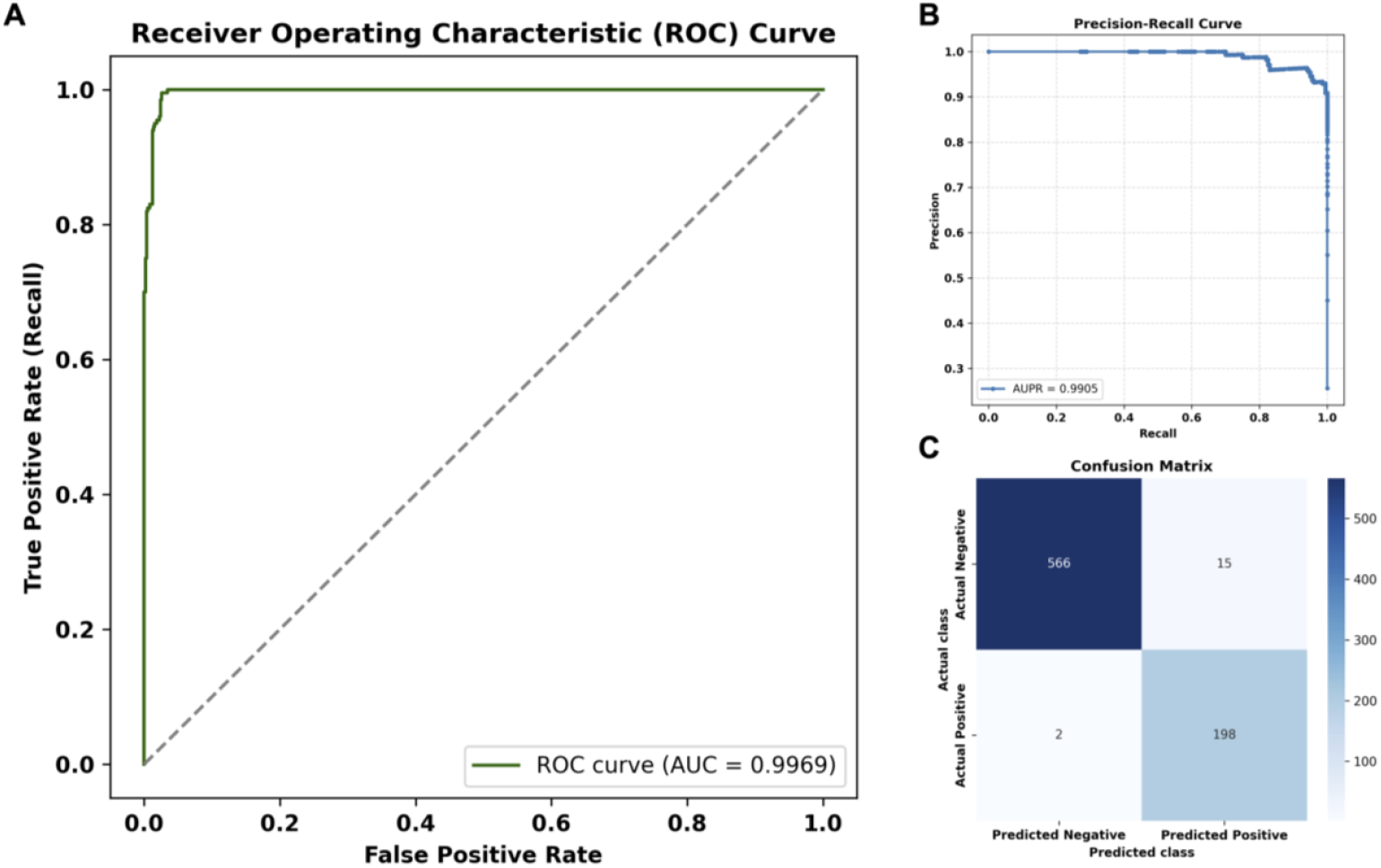
Performance of the best performing model for predicting the human infection potential of coronaviruses. (A) ROC curve showing the classification performance of the final model on the test dataset. (B) Precision-recall (PR) curve summarizing the trade-off between precision and recall, with the area under the PR curve (AUPR) shown. (C) Confusion matrix illustrating the number of true positives, true negatives, false positives, and false negatives obtained on the test dataset.

### Cross-Validated Reliability of the Optimized Random Forest Model

To assess its generalizability and stability, the performance of the Random Forest Classifier was validated using 10-fold cross-validation. The model achieved an average accuracy of 0.9736 ± 0.0071, demonstrating consistently high overall classification performance across folds. AUC was 0.9939 ± 0.0029, indicating excellent predictive ability. The mean F1 score was 0.9501 ± 0.0134, reflecting a strong balance between precision and recall. The AUPR averaged 0.9092 ± 0.0224, confirming the model’s robustness in handling class imbalance. The MCC was 0.9329 ± 0.0182, further supporting the high reliability and balanced predictive performance (Fig 4). The low standard deviations across all metrics demonstrate that the performance is stable and not overly dependent on any single fold of the dataset.

**Fig 4.**
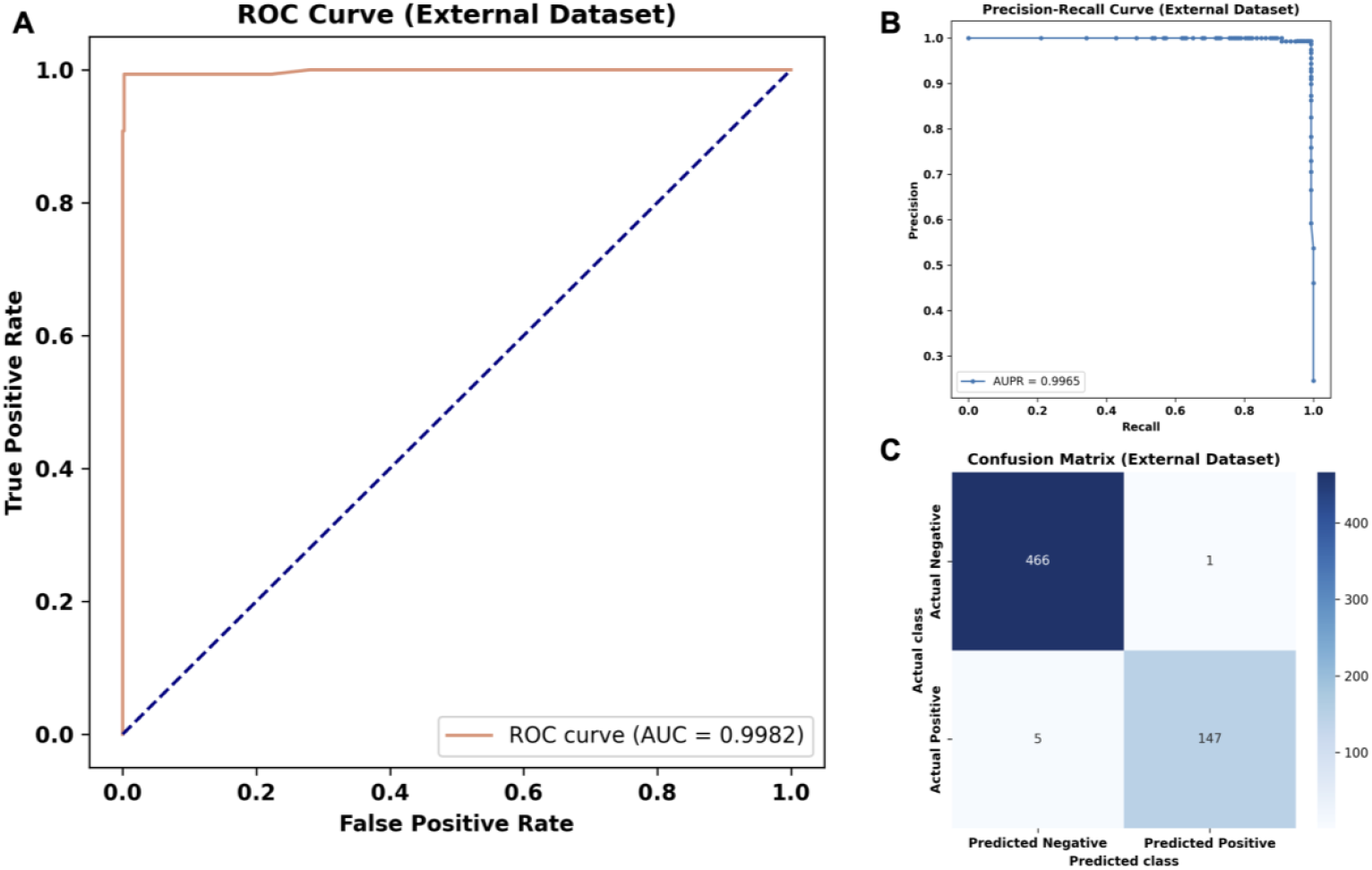
10-fold cross-validation performance of the Random Forest Classifier. (A) Distribution of performance metrics across k-fold cross-validation, shown as boxplots for accuracy, AUC, F1 score, AUPR, and MCC. Boxes indicate the interquartile range, center lines represent median values, and whiskers denote the full range of observed values. (B) Performance of each metric across individual cross-validation folds, illustrating model consistency and variability.

### External Validation Confirms Robust and Generalizable Model Performance

External validation demonstrated that the Random Forest Classifier maintained excellent predictive performance when applied to an independent dataset of coronavirus spike protein sequences. The consistently high scores across all key metrics including sensitivity, specificity, accuracy, AUC, F1 score, AUPR, and MCC of 0.9671, 0.9979, 0.9903, 0.9993, 0.9800, 0.9980, and 0.9738, respectively, indicated that the model generalizes well to unseen data and reliably predicts between human and non-human infection potential (Fig 5). These results confirmed the model’s robustness, high reliability, and practical utility for predicting the infection potential of newly emerging coronavirus.

**Fig 5.**
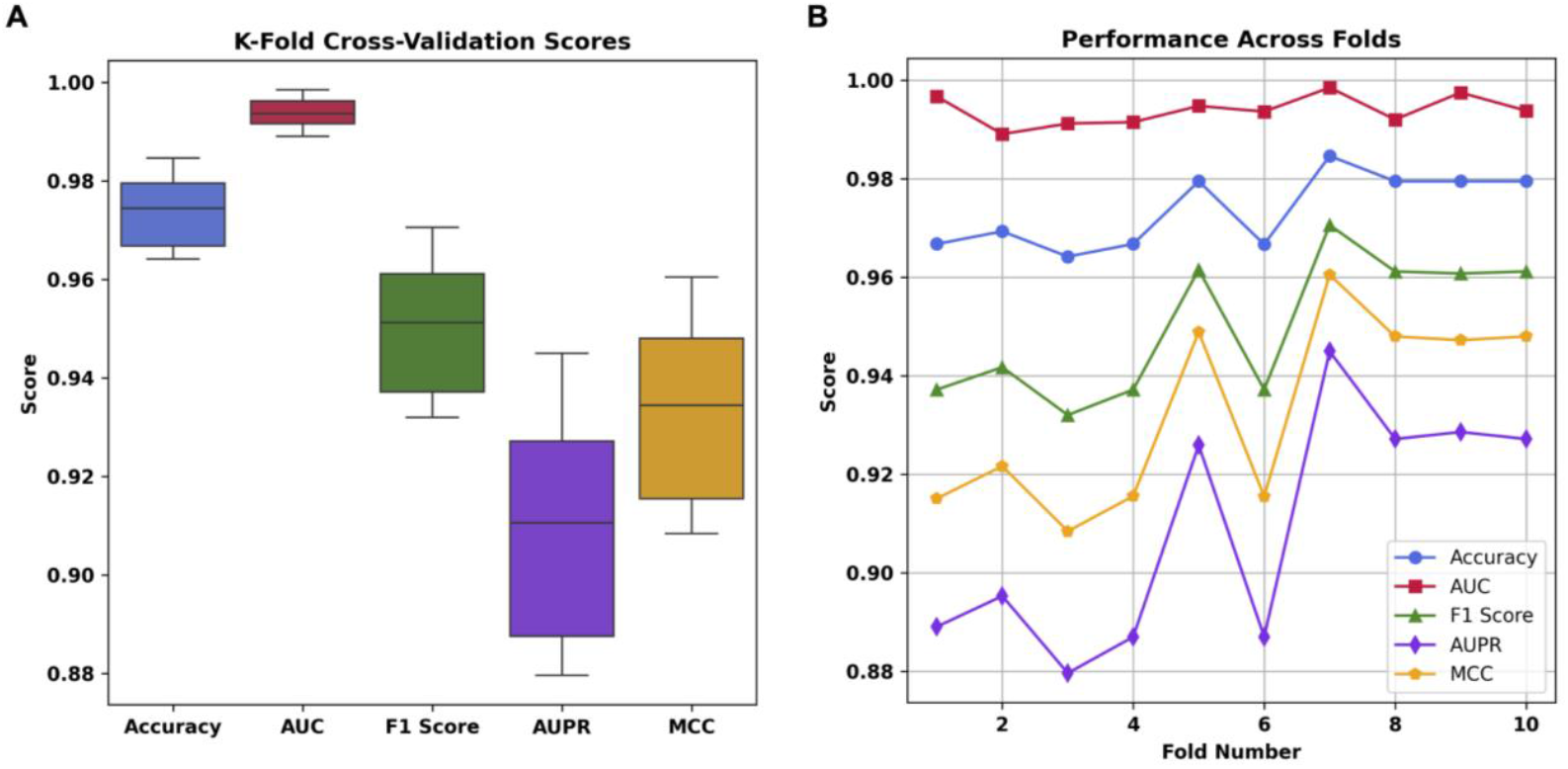
Performance of the Random Forest Classifier on the external dataset. (A) ROC curve showing the classification performance of the final model on the external dataset. (B) The PR curve summarizing the trade-off between precision and recall, with the AUPR shown. (C) Confusion matrix illustrating the number of true positives, true negatives, false positives, and false negatives obtained on the external dataset.

### SHAP-Based Identification of Associated Tri-Amino Acid Motifs

To identify tri-amino acid motifs potentially associated with human infection, SHAP analysis was applied. The tri-amino acid motif KIQ, LEP, TGS, NGD, TGE, YTA, and TVS demonstrated the top impact across all datasets, with elevated values of these features driving predictions toward the human infection and non-human infection group. Some features showed consistent directional effects on the model output, while others had variable influence depending on the sequence context (Fig 6A-G). These findings highlight specific sequence patterns that play a critical role in predicting human infection based on spike proteins and provide targets for further biological investigation.

**Fig 6.**
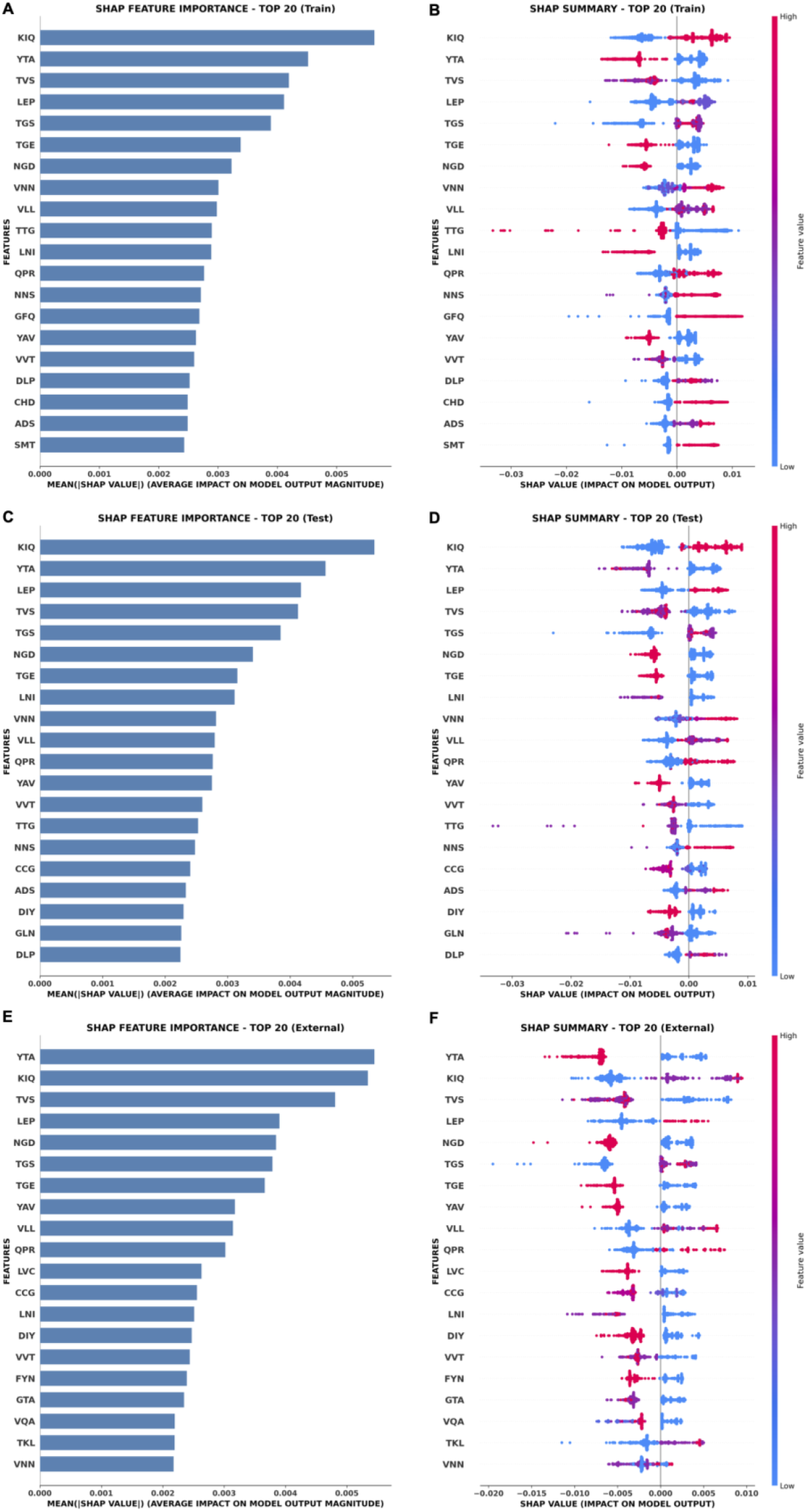
The top 20 most importance and impactful k-mer features for the Random Forest Classifier’s predictions. (A-C) Mean absolute SHAP values of features ranked by their average impact magnitude on the model’s output, regardless of direction. (D-F) SHAP value distribution. Each dot represents an individual sample’s SHAP value for a specific feature (k-mer). The position on the x-axis shows the impact of that feature on the model output (positive or negative influence on predicting human infection potential). The color represents the feature value (from low in blue and high in red).

### Motif Frequency Confirms Their Predictive Value for Human Infection

To confirm the association of the identified tri-amino acid motifs with human infection potential, the percentage occurrence of each motif was performed. Only KIQ and LEP were found at high frequencies in the human infection group (>70%), while remaining markedly have less frequent in the non-human infection group (<30%). In contrast, TGS was present at high frequencies in both human infection (98%) and non-human infection (48%) groups, indicating that it does not clearly discriminate between the two groups. Therefore, only KIQ and LEP were selected for further evaluation. Conversely, NGD, YTA, TVS, and TGE were enriched in the non-human infection group (>50%) but were low or absent in the human infection group (Fig 7A). Among these, only TVS reached a high frequency (>70%) in the non-human infection group. Collectively, these findings suggested that KIQ and LEP are positively associated with human infection potential, whereas NGD, YTA, TVS, and TGE are indicative of non-human infection potential.

**Fig 7.**
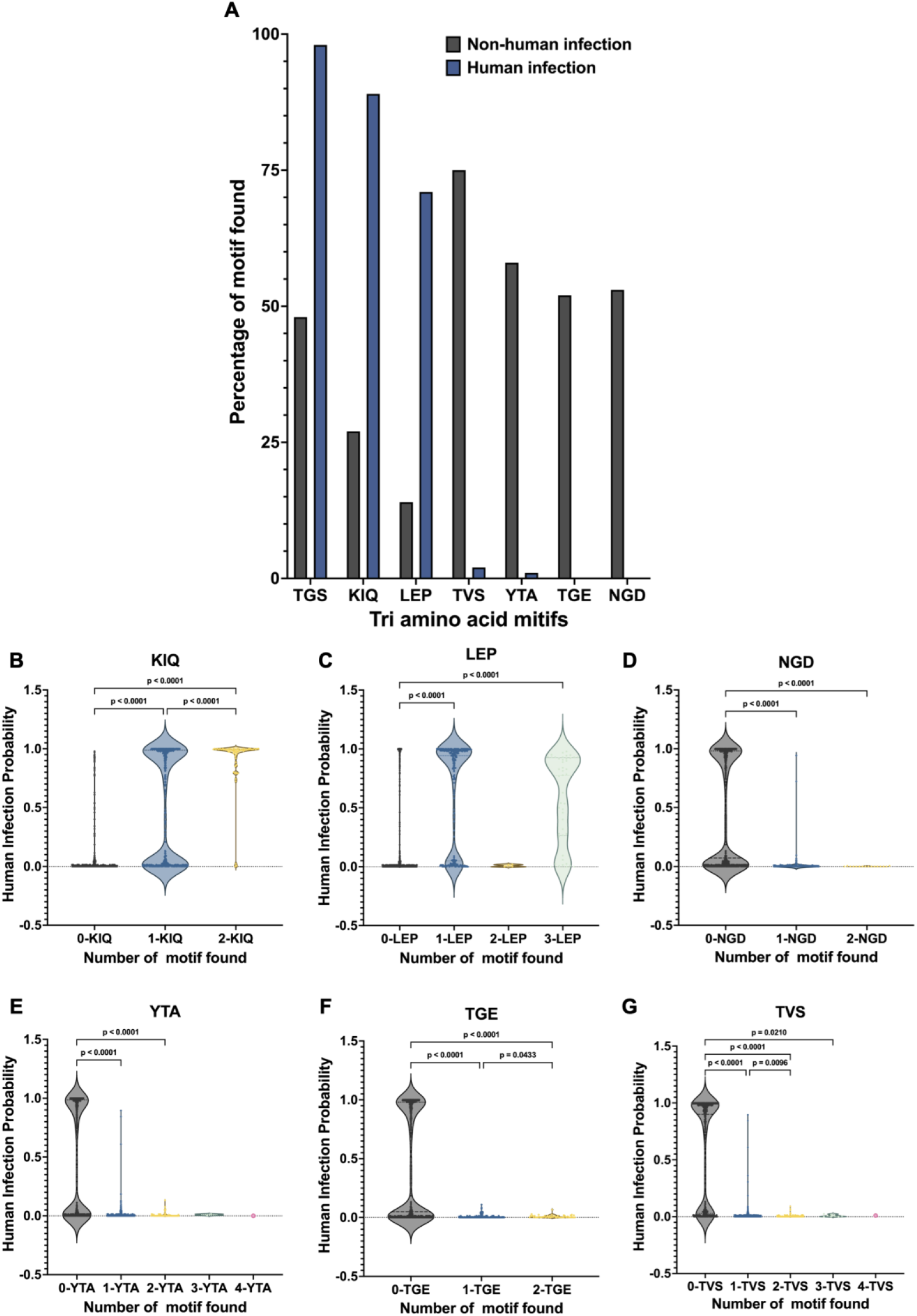
Confirmation of tri-amino acid motifs associated with human and non-human infection. (A) Bar plot showing the percentage occurrence of seven identified tri-amino acid motifs (KIQ, LEP, TGS, NGD, YTA, TVS, and TGE) across the complete dataset, including both original and external sequences. Grey bars represent the non-human infection group, and blue bars represent the human infection group. (B-G) The motif abundance relevant to human infection probability. The grey plots represent human infection probability associated with non-motif found, blue plots represent human infection probability associated with one motif found, yellow plots represent human infection probability associated with two motifs found, green plots represent human infection probability associated with three motifs found, and pink plots represent human infection probability associated with four motifs found.

To further evaluate the relationship between motif abundance and infection probability, the motif abundance relevant to human infection probability were plotted. For KIQ, an increased number of occurrences significantly corresponded to higher confidence in predicting human infection while LEP, significant increases in predicted human infection probability were observed only when one or three LEP motifs were present. In contrast, NGD, YTA, TGE, and TVS were significantly consistent linked to confident predictions of non-human infection when absent. (Fig 7B-G). Collectively, these results demonstrate that the frequency of specific tri-amino acid motifs corresponds with the probability of human infection.

### Localization of Key Tri-Amino Acid Motifs on Coronavirus Spike Proteins

To determine the location of the tri-amino acid motifs, KIQ and LEP were mapped onto the human infected SARS-CoV-2 spike protein sequence (Accession: YP_009724390.1) KIQ and LEP, which correspond to human infection potential, were found at positions 933-935 and 222-224 which located in the Heptad repeat 1 (HR1) and N-terminal domain (NTD) region, respectively (Fig 8A and 8C-D). In contrast, NGD, YTA, TGE, and TVS were mapped onto the spike protein of Infectious Bronchitis Virus (IBV) (Accession: AFJ11176.1), which has not been reported to infect humans. NGD and TVS were found in the NTD region at positions 169-171 and 148-150, respectively. YTA was found at positions 752-754 which located in the fusion protein (FP), while TGE was most commonly observed at positions 674-676 (Fig 8B and 8C-D).

**Fig 8.**
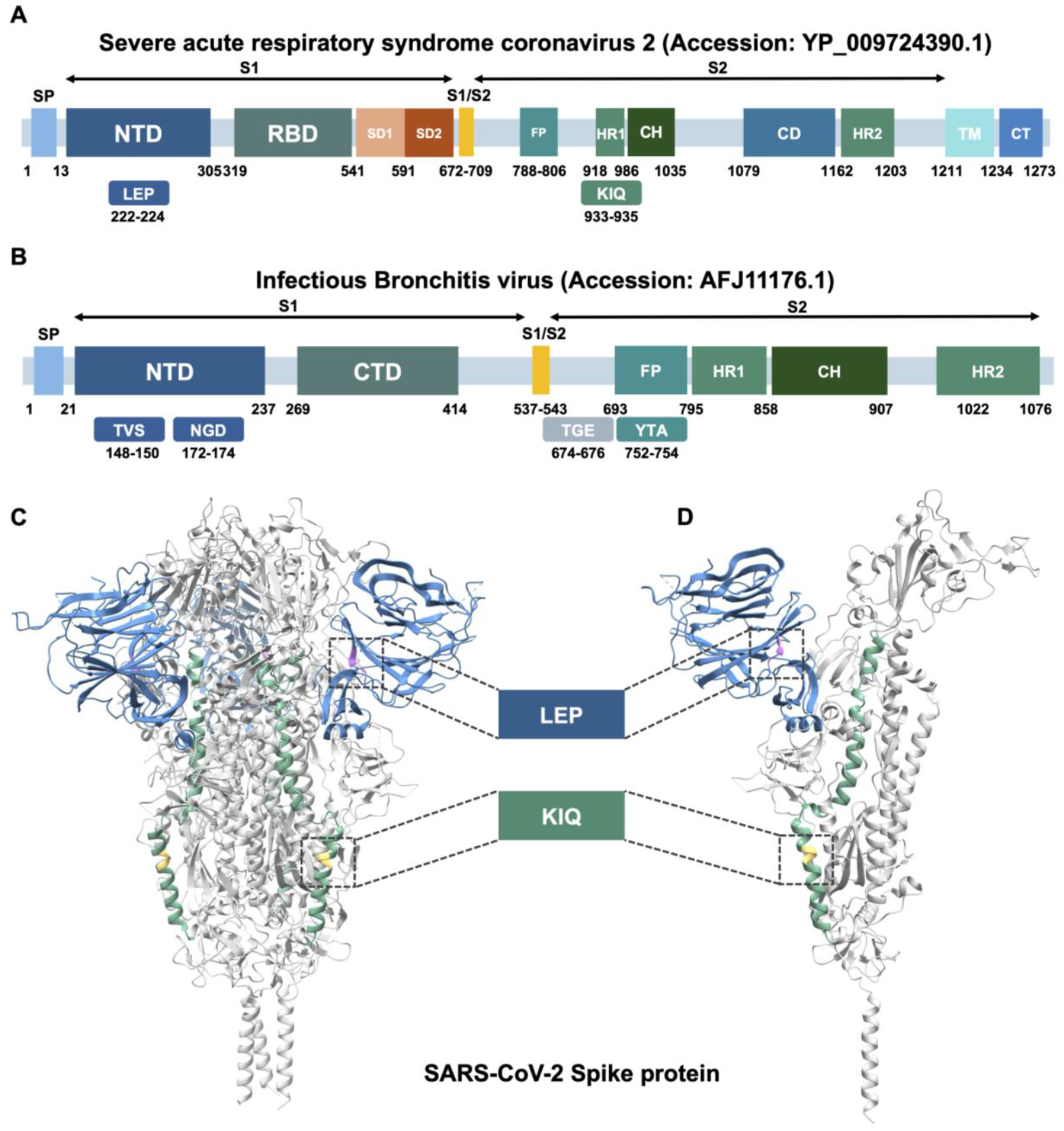
Mapping of identified tri-amino acid motifs onto spike protein sequences and structure. (A) SARS-CoV-2 spike protein sequence with motifs mapped to their positions. (B) Infectious bronchitis virus (IBV) spike protein sequence with motifs mapped to their positions. (C) Trimeric structure of the SARS-CoV-2 spike protein (PDB: 6XR8). (D) Monomeric structure of the SARS-CoV-2 spike protein. The N-terminal domain (NTD) is shown in blue, and the heptad repeat 1 (HR1) region is shown in green. The LEP motif is shown in purple, and the KIQ motif is shown in yellow.

## Discussion

Machine learning provides a fast and effective approach for screening large-scale cross-species pathogen transmission and assessing the human infection potential of various virus families. In this study, we developed machine learning-based framework to predict the human infection potential of coronaviruses based on spike protein sequences. Our dataset comprised 3,904 spike sequences representing the most comprehensive all genus level coverage reported to date including *alpha-, beta-, gamma-*, and *deltacoronavirus*. Moreover, we conducted systematic benchmarking of 27 machine learning models to identify the optimal predictive framework, ensuring robust model selection and generalizability. Beyond prediction, our workflow integrates feature interpretation to identify tri-amino acid motifs associated with human infection, providing biologically meaningful insights.

The Random Forest Classifier achieved the highest predictive performance across all tested machine learning models in our study with 97.8% accuracy, 99% sensitivity and 97.4% specificity. However, multiple machine learning algorithms also demonstrated consistently high accuracy, indicating that machine learning approaches are broadly effective and well suited for analyzing spike protein sequence-derived features. This contrasts with a previous study in which Logistic Regression was selected as the optimal classifier, achieving 99% accuracy, 99% sensitivity, and 99% specificity (23). A key methodological difference lies in the feature representation of protein sequences. The previous study used a skip-gram neural network model, which turns amino acid sequences into vector embeddings that capture patterns and relationships across the sequence. In contrast, our analysis utilized k-mer features, representing each sequence by the frequency of all possible trimers. While k-mer counting provides a transparent and interpretable feature space, it lacks the contextual embedding capabilities inherent in skip-gram models. Nevertheless, the superior performance of the Random Forest classifier in our framework suggests that k-mer based features, when paired with a robust ensemble algorithm, can effectively distinguish human infection from non-human infection coronaviruses. Moreover, the slightly lower performance reflects the increased sequence diversity and complexity resulting from the inclusion of the most comprehensive all genus of coronavirus, while previous studies achieved strong performance to detect the potential of coronaviruses to infect human but were limited to only *alpha-* or *betacoronavirus* (23, 24). Spike protein sequences differ markedly across genera in terms of length, domain organization, and amino acid composition, resulting in more heterogeneous and comprehensive sequence patterns.

In addition to predictive modeling, we identified tri-amino acid motifs associated with human and non-human infection potentials. Based on feature importance analysis, six key motifs were identified as highly influential for prediction. Among these, KIQ and LEP were enriched in human infection coronaviruses, while NGD, YTA, TGE, and TVS were more common in non-human infection viruses. These motifs differ from those reported in a previous study that developed a machine-learning pipeline for host prediction in the Coronaviridae family. In that work, the motif TLTN and IPQN in NTD and HR1 region, respectively, were identified as an important feature associated with human infection (25). The discrepancy between the motifs identified could be from the use of distinct feature-importance strategies. In their work, feature relevance was determined using Gini impurity reduction within a Random Forest framework, where tetra-mer sequence features contributing most to decreases in mean Gini impurity were prioritized. In contrast, we applied SHAP to quantify the contribution of tri-amino acid k-mer features to individual model predictions. Gini impurity is known to be biased toward high-frequency or high-cardinality features, Whereas SHAP provides a consistent, model-agnostic measure of feature contribution across the entire prediction space. As a result, our approach highlights motifs that exert stable and interpretable effects on human infection potential across diverse coronavirus genera, whereas impurity-based methods may emphasize different, region- or frequency-driven sequence patterns.

The KIQ motif, located at positions 933-935 within the HR1 region, lies in a domain essential for SARS-CoV-2 membrane fusion. After exposure and cleavage of the S1/S2 site by host proteases such as TMPRSS2, the FP protein inserts into the host membrane, initiating formation of the HR1-HR2 six-helix bundle. HR1 undergoes a major “jack-knife” refolding that extends the central coiled-coil structure and exposes hydrophobic surface grooves. Heptad repeat 2 (HR2) subsequently binds these grooves in an antiparallel manner to assemble the six-helix bundle that drives membrane fusion (26). Importantly, KIQ and adjacent residues have been reported as highly conserved spike-specific CD4^+^ T-cell epitopes capable of eliciting SARS-CoV-2 S-specific CD4^+^ and CD8^+^ memory T cells following COVID-19 vaccination (27). This suggests that the KIQ motif may play a crucial role in SARS-CoV-2 fusion and vaccine-induced immunity.

Notably, LEP was located at positions 222-224 within the NTD of the SARS-CoV-2 spike protein which know as an important binding region to interact with the tyrosine-protein kinase receptor UFO (AXL), facilitating infection of pulmonary and bronchial epithelial cells (28, 29). Previous structural studies identified residues Lys147-Lys150, Trp152, Arg246, Ser247, and Ser256 of the NTD as interacting with AXL residues Pro57, Pro58, Glu59, His61, Ile68, Glu70, Glu85, and Phe113 (28). Taken together, these findings indicate that the identified motifs, particularly LEP and KIQ, may have functional relevance in receptor engagement, membrane fusion, and host adaptation. Nonetheless, their precise biological roles require further investigation through targeted *in vitro* assays and mutagenesis experiments.

Our dataset was annotated into two categories including human infection and non-human infection based on the reported host species from which each viral sequence was isolated. This classification relies on available host data in NCBI virus databases rather than experimental evidence of host range. Therefore, it should be noted that some sequences categorized as “non-human infection” have not been experimentally confirmed as incapable of infecting humans. This introduces a potential source of uncertainty in our dataset and represents one of the limitations of our study.

Overall, these findings highlight the value of integrates systematic model benchmarking, robust performance evaluation, and local model interpretability with sequence-level mapping to not only build accurate predictive tools but also generate biologically meaningful insights. This integrated approach provides a practical framework for the risk assessment of novel emerging coronaviruses and could support future surveillance and preparedness efforts to mitigate potential cross-species transmission.

## Acknowledgement

This research was supported by the Faculty of Medicine, Khon Kaen University, Thailand (Grant No. IN68103), the European Union’s Horizon Europe Research and Innovation Programme (Grant Agreement No. 101095444, PANDASIA), and the Volkswagen Foundation (Project No. 9C 451, PANDA).

## Supplementary data

All data supporting the results of this study are provided in the Supplementary Information.

**S1 Table. Original dataset of spike protein sequence**. Complete list of coronavirus spike protein sequences used in this study, including virus name, coronavirus genus, accession number, amino acid sequence, and human infection annotation. This dataset represents all four coronavirus genera and served as the primary training and testing dataset.

**S2 Table. External dataset of spike protein sequence with human infection probability**. External coronavirus spike protein sequences including virus name, coronavirus genus, accession number, amino acid sequence, human infection annotation, and predicted probabilities of human infection generated by the final machine learning framework. This dataset was used to evaluate model generalizability and evaluate the motif abundance relevant to human infection.

**S3 Table. Model benchmarking results**. Performance metrics of all machine learning models evaluated in this study across 14 repeated runs. This table summarizes the comparative performance of each model for predicting the human infection potential of coronaviruses.

**S4 Table. Summary of model benchmarking (mean ± SD)**. Summary statistics of model performance across 14 repeated runs, reported as mean ± standard deviation. This table highlights model robustness and stability and was used to guide selection of the optimal predictive model.

**S5 Table. Hyperparameter tunning results**. Results of hyperparameter optimization for the selected machine learning models, including parameter ranges tested and corresponding performance outcomes. These results informed the final model configuration used for downstream prediction and interpretation analyses.

